# Variant emergence, not vaccine deployment, drives episodic positive selection on the SARS-CoV-2 spike at provincial scale in Canada

**DOI:** 10.64898/2026.05.16.725625

**Authors:** Marissa Tomas, Pape Cheikh Khary Ndongo, Matthieu Vilain, Stéphane Aris-Brosou

## Abstract

Mass immunization against SARS-CoV-2 created a heterogeneous landscape of antibody-mediated immune pressure, yet whether this pressure measurably altered episodic positive selection on spike remains unresolved. Using Canadian genomic surveillance data spanning the five major variants of concern (Alpha, Beta, Gamma, Delta, and Omicron), we inferred time-resolved phylogenies from spike-coding sequences and applied site- and branch-level episodic selection models to identify when and where adaptive change occurred. To evaluate whether vaccination intensity was associated with selection, we integrated these phylogenetic analyses with provincial vaccination time series using cross-correlation and lagged panel regression models that accounted for province and time effects, lineage prevalence, and sampling heterogeneity. Episodic positive selection was concentrated at a limited number of spike codons, especially within the N-terminal domain, receptor-binding domain, and furin cleavage region. However, these signals were dominated by substitutions associated with variant emergence, particularly during the Alpha-to-Delta transition, rather than by vaccination rollout. Whole-gene tests provided no evidence that vaccine intensity was associated with elevated episodic selection, and residualized vaccination trajectories did not predict selection at biologically plausible lags. Across provinces, the timing and distribution of selection events were inconsistent with a vaccine-driven escape model. Together, these results indicate that, at provincial resolution in Canada, episodic positive selection on SARS-CoV-2 spike was driven primarily by variant turnover rather than vaccine deployment. More broadly, this study provides a quantitative, VOC-resolved assessment of spike evolution in a structured epidemic and suggests that population-level vaccination intensity was not a detectable determinant of spike adaptation in the period examined.

## Introduction

The SARS-CoV-2 pandemic, caused by a betacoronavirus first identified in Wuhan, China in late 2019, resulted in over 700 million confirmed infections and more than six million deaths worldwide before endemic transition (World Health Organization, 2024). The homotrimeric surface-exposed spike (S) glycoprotein is the principal determinant of viral tropism, mediating host cell entry through binding of the angiotensin-converting enzyme 2 (ACE2) receptor via its receptor-binding domain (RBD), and constitutes the near-exclusive target of potently neutralizing antibodies elicited by both natural infection and vaccination (Walls et al., 2020; Wrapp et al., 2020). This dual functional role acts as both an entry machinery and an immunodominant antigen, hereby placing the spike protein at the intersection of viral fitness and immune recognition, and making it the primary substrate for adaptive evolution under mounting population-level immune pressure.

Following the global spread of the ancestral Wuhan-Hu-1 lineage, SARS-CoV-2 underwent rapid and repeated phenotypic diversification, giving rise to a succession of variants of concern (VOCs): Alpha (B.1.1.7), Beta (B.1.351), Gamma (P.1), Delta (B.1.617.2), and Omicron (B.1.1.529) (Lauring and Hodcroft, 2021; Carabelli et al., 2023). Each VOC had distinctive constellations of mutations in the spike protein, concentrated in the RBD and the N-terminal domain (NTD), that collectively altered transmissibility, cell tropism, and, critically, susceptibility to neutralization by infection- and vaccine-induced antibodies (Greaney et al., 2021; Planas et al., 2021). The recurrent, convergent emergence of functionally equivalent mutations across independently evolving lineages, such as E484K/E484A at a key ACE2-contact and antibody-epitope site, acquired independently in Beta, Gamma, and Omicron, represents compelling phenotypic evidence that antigenic escape from pre-existing immunity constitutes a persistent selective force acting on spike (Obermeyer et al., 2022; Rochman et al., 2021).

Formal phylogenetic tests based on codon-substitution models have provided a statistically principled framework for quantifying these selective pressures (Yang, 2007). Site-level approaches, notably the Mixed Effects Model of Evolution (MEME; Murrell et al., 2012) implemented in HyPhy (Pond, Frost and Muse, 2005; Weaver et al., 2018), have identified numerous spike codons evolving under episodic positive selection across VOC lineages. However, site-level models are inherently agnostic to the temporal and phylogenetic context in which selection intensifies: they cannot resolve which lineages, and at what point in the epidemic, selective pressures were elevated. This limits their utility for testing epidemiological hypotheses about the drivers of viral adaptation. A more powerful alternative that directly addresses this limitation is the Branch-Site Unrestricted Test for Episodic Diversification (BUSTED; Murrell et al., 2015), also implemented in HyPhy. Crucially, BUSTED permits the designation of *foreground* branches (those hypothesized to be under elevated selection) against a *background* in which *ω* ≤ 1 is enforced, and evaluates, via a likelihood ratio test, whether there is gene-wide evidence for positive selection (*ω >* 1 in at least a proportion of codons) specifically on those foreground lineages. This foreground-background partitioning maps directly onto our epidemiological design: lineages circulating in provinces and time windows characterized by high vaccine coverage constitute the foreground, while those predating or occurring under minimal immune pressure serve as the background. Rather than requiring a two-step procedure in which selected branches are first identified and then correlated with vaccination data post hoc, BUSTED embeds the epidemiological hypothesis within the phylogenetic model itself, yielding a single, direct test of whether vaccine-derived immune pressure is associated with gene-wide adaptive evolution of the spike protein. This framing is also biologically appropriate: immune evasion of the spike does not typically operate through a single codon but through coordinated mutational changes across the RBD and NTD (Greaney et al., 2021; Carabelli et al., 2023), making a gene-wide test better matched to the underlying biology than a site-by-site decomposition.

Mass immunization campaigns, deployed across Canada and globally from December 2020 onward, introduced a spatially and temporally heterogeneous source of antibody-mediated immune pressure onto circulating SARS-CoV-2 populations that was qualitatively distinct from that of prior infection. There is growing consensus that humoral immunity, from infection, vaccination, or hybrid exposure, is now the dominant selective force shaping the evolutionary trajectory of the virus (Kistler, Huddleston and Bedford, 2022; Raharinirina et al., 2025). Canadian provinces differed substantially in both the pace and coverage of vaccine rollout due to logistical, demographic, and policy factors (Fawcett, 2021; Fitzpatrick et al., 2023), creating a natural quasi-experimental framework in which the intensity of vaccine-derived immune pressure on co-circulating viral lineages varied across space and time. Whether this heterogeneity translates into measurable differences in the intensity or timing of positive selection on the spike protein, and whether such differences are consistent across the five major VOCs, has not been formally evaluated.

Here, we address this question by integrating foreground-partitioned phylogenetic selection analyses with provincial vaccination surveillance data across Canada. We apply MEME and BUSTED analyses to curated spike-coding alignments from the five major VOCs sequenced in Canada and deposited in the GISAID database (Shu and McCauley, 2017), designating as foreground those lineages associated with provinces and time intervals of elevated vaccine coverage, and test for gene-wide signatures of episodic positive selection on those foreground branches. Temporal dependencies between vaccination intensity and selection signals are further modeled through cross-correlation analysis and lagged panel regression with province and time fixed effects, providing estimates of the magnitude and direction of these associations while controlling for confounding factors including lineage prevalence and sampling effort. By embedding the epidemiological hypothesis directly within the phylogenetic model, this study contributes quantitative evidence to the emerging framework in which vaccination campaigns are recognized as agents of viral evolution, with direct implications for the timing and antigenic composition of future vaccine formulations.

## Materials and methods

### Genetic data retrieval and processing

All available SARS-CoV-2 genomic sequences were downloaded, along with their metadata, from the GISAID database (Shu and McCauley, 2017) on December 19, 2025. Next, sequences were filtered based on their metadata, keeping only those marked as complete, with high coverage, a complete collection date, and collected in Canada. Then, the sequences were aligned to the SARS-CoV-2 reference sequence NC_045512.2 available on the National Center for Biotechnology Information (National Center for Biotechnology Information, 2020). The alignment was performed using MAFFT ver.7.526 (Katoh and Standley, 2013) with the “–6merpair –thread 8 –keeplength –addfragments” options. After alignment, the Open Reading Frame (ORF) of the spike protein was extracted, and sequences that did not start with a methionine codon (ATG) or that did not end with a stop codon (TAA, TAG, or TGA), and that thus were out of frame, were removed. The final preprocessing step consisted of cleaning the alignment to remove low-quality or poorly aligned sequences. To do so, the overlap trimming method described in the trimal ver.1.5.1 software (Capella-Gutiérrez, Silla-Martínez and Gabaldón, 2009), which currently is not suited for large alignments, was reimplemented in Python. This fork defines an overlap as having a gap in both positions, an indetermination (*N*) in both positions, or a residue in both positions. For each sequence at each position in the alignment, the percentage of overlap with all other sequences was computed. Finally, good positions were defined as overlapping with at least 1% of all sequences, and we excluded all sequences with less than 99% of good positions.

The codon alignment was then split by province, making sure each dataset has a copy of the reference sequence (NC_045512.2), and filtered to remove sequences containing indels (as obvious sequencing errors). Ambiguous nucleotide were imputed based on majority rule consensus, and stop codons were removed. Tip dates were parsed from the `YYYY_MM_DD` suffix of each sequence identifier. In order to make said analyses computationally manageable, the sequences were then filtered with CD-HIT (Fu et al., 2012) at the 100% similarity level. A maximum phylogenetic tree was the inferred for each province in R ver.4.5.1 (Hornik, 2012), with phangorn ver.2.12.1 (Schliep, 2011), under the HKY model of evolution (Hasegawa, Kishino and Yano, 1985) +Γ_4_ (Yang, 1994). Each tree was rerooted with the reference sequence, and a script was written to create alignment files containing the estimated trees for the selection analyses with HyPhy ver.2.5.78 (Pond, Frost and Muse, 2005; Kosakovsky Pond et al., 2020).

### Selection analyses

HyPhy requires a stop-codon-free codon alignment with no ambiguous bases in interpreted reading frames (Kosakovsky Pond et al., 2020). For each provincial alignment, we therefore replaced every premature stop codon (TAA, TAG, TGA) with the most frequent non-stop codon at the same alignment column. Single-nucleotide IUPAC ambiguity codes were resolved column-by-column to the most frequent IUPAC-compatible base; positions that remained polymorphic across the column were assigned the column consensus. The two operations were performed in a single pass, with all substitutions recorded for provenance. For instance, the British Columbia alignment underwent 4,034 ambiguity imputations and one stop-codon replacement (TAG → GAG at codon 1,262).

### Bayesian molecular-clock inference

A Bayesian time-calibrated phylogeny was inferred for each province using BEAST v1.10.5 (Suchard et al., 2018). Substitution was modeled with HKY+Γ_4_ (Hasegawa, Kishino and Yano, 1985; Yang, 1994); molecular evolution was assumed to follow a strict molecular clock; and the coalescent demographic prior was set to a constant population size. Tip dates were fixed to the collection day of each sample, allowing the clock rate to be jointly inferred from temporal sampling structure. For each province, two independent Markov chain Monte Carlo (MCMC) sampler were run for 10^8^ states each, with parameter and tree samples logged every 10^4^ states. The chains were combined with logcombiner v1.10.5 after discarding the first 5 × 10^7^ states of each chain (50% burn-in). The maximum-clade-credibility (MCC) tree was summarized with treeannotator v1.10.5 using mean node heights.

### Convergence diagnostics

Per-parameter effective sample size (ESS) was computed with the initial monotone sequence estimator of (Geyer, 1992) on the post-burn-in samples of each chain, and the Gelman-Rubin potential scale-reduction statistic (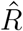; (Gelman and Rubin, 1992)) was computed across the two chains. A run was considered acceptably mixed when ESS ≥ 200 for every continuous parameter and 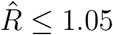 between the two chains, in line with the recommendations of (Rambaut et al., 2018). Chains failing either criterion (here, the New Brunswick and the Prince Edward Island datasets because of their small sample size) were flagged and their downstream estimates are reported only with explicit caveats.

### Cross-provincial posterior comparison

Marginal posterior distributions for the clock rate, root height, constant coalescent population size, transition/transversion ratio (*κ*), Gamma shape (*α*) and total tree length were compared across provinces by extracting the post-burn-in samples for each parameter and computing the median, mean, and the Hight Posterior Density (HPD) with DendroPy v4.6 (Sukumaran and Holder, 2010); results are summarized in SI file `compare_provinces.xlsx`.

### Episodic positive-selection inference

Site-wise tests for episodic diversifying selection were performed with MEME (Murrell et al., 2012) as implemented in HyPhy v2.5 (Kosakovsky Pond et al., 2020). A maximum-likelihood phylogeny under HKY+Gamma_4_ was first inferred from the cleaned codon alignment to provide a fixed input topology to HyPhy. MEME was run with default settings, returning per-site likelihood-ratio statistics together with the posterior probability that each branch belonged to the positive-selection *ω* class. Whole-gene episodic selection was tested with BUSTED (Murrell et al., 2015).

For each MEME hit, an empirical Bayes factor (EBF) was computed for every internal and terminal branch as

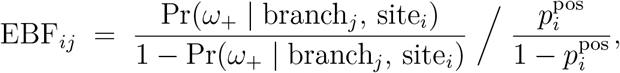

where the numerator is the branch-conditional posterior odds of the positive-selection class returned by MEME and the denominator is the prior odds derived from the site-level mixing weight 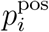 (i.e., the site-level mixture weight on the positive selection *ω* class). A (site × branch) event was retained as a *strict* episodic selection event when both *P*_MEME_ ≤ 0.10 and EBF ≥ 100, and as *suggestive* when *P*_MEME_ ≤ 0.20 and EBF ≥ 100.

### Dating selection events on the molecular-clock tree

Because HyPhy and BEAST inferred independent topologies on the same provincial dataset, branches were matched between the two trees by their descendant tip-set rather than by branch label. Tip names were normalized, to its embedded format `YYYY_MM_DD` showing collection dates. Each HyPhy branch was then identified with the unique BEAST branch sharing the same set of normalized descendant tips. Branches without a tip-set match (typically due to small topological discordance near short internal nodes) were excluded from dating; the per-province match rate ranged from 94% (ab) to 97% (nl). For every matched event, the parent and child node ages on the BEAST MCC tree were converted to calendar dates, and their HPD were retained. The calendar date attributed to each selection event was the child-node age, with the parent-child interval giving the temporal window over which the substitution causing the inferred event must have occurred.

### Provincial vaccine-deployment timeline

A long-format timeline of Canadian COVID-19 vaccine deployment events was assembled covering: (i) Health Canada authorization dates for the five primary-series vaccines (Pfizer-BioNTech BNT162b2, Moderna mRNA-1273, AstraZeneca/Covishield ChAdOx1-S, Janssen Ad26.COV2.S, Novavax NVX-CoV2373); the three bivalent boosters (Moderna BA.1 mRNA-1273.214, Pfizer BA.4/5 BNT162b2-Bivalent, Moderna BA.4/5 mRNA-1273.222); and the two XBB.1.5-adapted monovalent updates; (ii) provincial first-dose dates collated from provincial press releases and contemporaneous national reporting; and (iii) one-dose population-coverage milestones at 50% and 75% (Public Health Agency of Canada, 2024a). Per-province weekly counts of new first-dose administrations were obtained from the COVID-19 Canada Open Data Working Group’s `vaccine_administration_dose_1_pt` release (Berry et al., 2023), which redistributes the daily values published by the Public Health Agency of Canada through the *Health Infobase* portal (Public Health Agency of Canada, 2024b). Daily counts were clipped at zero (to absorb the rare retrospective downward corrections that appear in provincial source files) and aggregated to Monday-anchored weeks; for figure-level reporting, weekly counts were normalized to events per 100,000 residents using the 2021 mid-year provincial population estimates of Statistics Canada (Statistics Canada, 2021).

### Variant-phase covariate

A national variant-of-concern (VOC) phase indicator was constructed by assigning each calendar week to one of nine successive Canadian dominance phases: B.1 wildtype (until 2021-03-01), Alpha (2021-03-01; 2021-07-01), Delta (2021-07-01; 2021-12-15), Omicron BA.1 (2021-12-15; 2022-03-01), BA.2 (2022-03-01; 2022-07-01), BA.5 (2022-07-01; 2022-11-01), post-BA.5 (2022-11-01; 2023-04-01), XBB (2023-04-01; 2023-11-01) and JN.1 (from 2023-11-01). Phase boundaries were set at the calendar week in which the focal lineage first exceeded 50% of sequenced isolates in the integrated PHAC clinical and wastewater surveillance system (Berry et al., 2024), with cross-checks against the National Collaborating Centre for Infectious Diseases variant updates (National Collaborating Centre for Infectious Diseases, 2024). Pango lineage assignments follow previous work (Rambaut et al., 2020; O’Toole et al., 2022).

### Detrended cross-correlation analysis

Per-province monthly and weekly time series were constructed for the strict episodic selection event count and the dose-1 deployment series. Both series share variant emergence as a common driver: each new VOC increases the rate at which defining substitutions are fixed onto branches of the genealogy, and simultaneously raises the intensity of provincial genomic surveillance, which controls the number of branches available for site-by-branch inference, while each new VOC also triggers a public-health response that includes a vaccine campaign or a booster recommendation. The raw Pearson correlation between two such co-trending series is dominated by their common trend in the classical spurious-regression sense (Yule, 1926; Granger and Newbold, 1974) and would not distinguish a genuine vaccine-driven immune-escape signal from the unrelated co-occurrence of variant waves and vaccine campaigns. To recover the within-variant-phase component of the coupling, we therefore residualized both series against the national variant-phase covariate of Section *Variant-phase covariate* before computing the cross-correlation function (CCF). We note that this choice deliberately discards any component of vaccine-driven selection that is itself collinear with variant phase (because boosters are scheduled in response to variant emergence, the two are partly aligned by design), and so the analysis is statistically conservative against the vaccine-pressure hypothesis. We judged the alternative (leaving the shared trend in) to be a worse failure mode for a population-scale inference of causal direction.

Specifically, with **y** ∈ ℝ^*T*^ a province-level series (selection or vaccine), **X** ∈ {0, 1}^*T* ×*K*^ the one-hot phase design matrix and 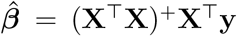 the ordinary-least-squares estimate via the Moore-Penrose pseudoinverse, the residual series 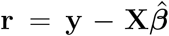 was retained for cross-correlation analysis. The Pearson product-moment correlation between residualized selection and residualized vaccine deployment was computed at integer lags *τ* ∈ {−8, …, +24} weeks, with the convention that positive *τ* implies that selection follows vaccine deployment by *τ* weeks. A 1,000-replicate circular-shift permutation test was used to assess significance: at each replicate the residualized vaccine series was rolled by a random integer drawn from {8, …, *T* − 8}, the cross-correlation function recomputed and its maximum recorded; the empirical *p*-value of the observed maximum is reported as `perm-p`. The 95th percentile of the null max-correlation distribution is shown as a horizontal threshold in each per-province panel of the cross-correlation figure.

### Visualization, validation and code availability

All numerical analyses were carried out in R ver.4.5.1 (Hornik, 2012) and in Python 3 using numpy (Harris et al., 2020), pandas (The pandas development team, 2020), scipy (Virtanen et al., 2020), numba (Lam, Pitrou and Seibert, 2015) and DendroPy (Sukumaran and Holder, 2010).

In order to confirm the validity, verifiability and reproducibility of all analyses, three versions of all the scripts developed in this work were written, coding both manually (by PCKN), and by vide-coding (Pimenova et al., 2025) under two models: Claude’s Sonnet 4.6 (by MT) and Claude’s Opus 4.7 (by SAB).

Vector-format figures were produced with matplotlib (Hunter, 2007) and exported as PDF. Reproducibility scripts (the aligner, the per-province BEAST dispatcher and their xml files, convergence diagnostics, the selection-event dating pipeline, the vaccine-timeline builder and the cross-correlation analyses; as described in the Supplementary material) are available from https://github.com/sarisbro/data.

## Results

### Bayesian clocks recover 8/10 credible time-calibrated trees

Provincial BEAST analyses produced time-calibrated phylogenies whose estimated clock rate, root height, transition/transversion ratio, Γ shape parameter, total tree length and constant coalescent population size are summarized in Table 1. Across the nine provinces that converged, the inferred substitution rate of the spike coding sequence ranged from 7.3 × 10^−4^ substitutions/site/year in Newfoundland & Labrador to 3.0 × 10^−3^ substitutions/site/year in British Columbia, with most provinces clustering between 1.7 × 10^−3^ and 2.2 × 10^−3^ substitutions/site/year. These values are within the published range for SARS-CoV-2 (Suchard et al., 2018) and agree to within the HPD of any pair of pandemicera estimates. The most recent common ancestor of each provincial tree dated to mid-to late 2019 in every converged province. The British Columbia and Saskatchewan trees, in particular, recover root dates compatible with the early November 2019 introduction window inferred from global SARS-CoV-2 genomic epidemiology. The Prince Edward Island analyses failed to mix in either MCMC chain (joint-likelihood ESS *<* 20, with posterior mass for the root height collapsing onto values orders of magnitude beyond the calendar window of pandemic sampling), were deemed unreliable, and discarded from downstream analyses. Newfoundland and Labrador and Nova Scotia each mixed acceptably but their HPD for the root height span several years, reflecting their small sample sizes (21 and 67 tips, respectively).

**Table 1.** Per-province posterior summaries from the BEAST molecular-clock analyses. Values are posterior means with 95% highest-posterior-density (HPD) intervals in parentheses, estimated from the combined post-burn-in samples of two independent MCMC chains. Sample size *n* is the number of tip sequences retained in the cleaned codon alignment. The whole-gene BUSTED test for episodic positive selection (Murrell et al., 2015) is reported as *P*_*BUSTED*_.

| Province | $n$ | Clock rate ($\times 10^{-3}$ s/s/yr) | Root height (yr) | Root date (calendar) | Strict events | $P_{BUSTED}$ |
| --- | --- | --- | --- | --- | --- | --- |
| Alberta (AB) | 576 | 2.05 (1.76, 2.36) | 2.62 (2.32, 3.36) | 2019.61 (2019.31, 2019.78) | 46 | 0.31 |
| British Columbia (BC) | 1330 | 2.97 (2.68, 3.26) | 2.01 (1.94, 2.14) | 2019.84 (2019.78, 2019.92) | 185 | 0.20 |
| Manitoba (MB) | 152 | 2.15 (1.66, 2.66) | 1.79 (1.62, 2.01) | 2019.74 (2019.53, 2019.92) | 5 | 0.50 |
| New Brunswick (NB) | 29 | 1.29 (0.43, 2.30) | 2.10 (1.46, 3.72) | 2019.27 (2017.66, 2019.91) | 0 | 0.48 |
| Newfoundland & Labrador (NL) | 21 | 0.73 (0.22, 1.45) | 3.93 (1.43, 7.41) | 2017.42 (2013.93, 2019.91) | 0 | 0.50 |
| Nova Scotia (NS) | 67 | 0.95 (0.60, 1.33) | 3.06 (1.76, 4.57) | 2018.41 (2016.91, 2019.71) | 0 | 0.50 |
| Ontario (ON) | 588 | 1.66 (1.46, 1.86) | 4.45 (4.17, 4.77) | 2019.48 (2019.15, 2019.75) | 53 | 0.05 |
| Prince Edward Island (PE)<sup>†</sup> | 10 | 1.11 (0.00, 2.17) | — | — | 0 | 0.33 |
| Quebec (QC) | 1121 | 2.15 (1.94, 2.37) | 2.30 (2.12, 2.52) | 2019.60 (2019.39, 2019.79) | 179 | 0.28 |
| Saskatchewan (SK) | 168 | 2.17 (1.65, 2.71) | 2.24 (2.06, 2.47) | 2019.73 (2019.50, 2019.91) | 12 | 0.46 |

<sup>†</sup> Prince Edward Island did not converge: BEAST joint-likelihood ESS < 20 across both chains, $\hat{R} > 1.05$ for the root height and clock rate, and the root-height posterior collapses onto biologically implausible values. PE event totals and any per-event dates from this analysis are reported only for transparency and should not be interpreted.

### Episodic positive selection clusters on a small set of variant-defining spike sites

HyPhy MEME identified 480 strict (site × branch) episodic selection events (*P*_*MEME*_ ≤ 0.10, EBF ≥ 100) and 812 suggestive events (*P*_*MEME*_ ≤ 0.20, EBF ≥ 100) across the ten provinces. Strict events were concentrated in the four provinces with the largest sequence sets and were vanishingly rare or absent in provinces with small sample sizes: British Columbia (*n* = 185 events), Quebec (*n* = 179), Ontario (*n* = 53), Alberta (*n* = 46), Saskatchewan (*n* = 12), Manitoba (*n* = 5), and Nova Scotia, Newfoundland and Labrador, New Brunswick and Prince Edward Island (no strict hits; Table 1). The whole-gene BUSTED test was non-significant in every province; only Ontario approached near significance (*P*_*BUSTED*_ = 0.051), so the overall picture is one of narrow site-level signal rather than gene-wide diversifying selection.

The cross-province distribution of those site-level events is heavily non-uniform (Figure 1a). Three codons harbor strict hits in five different provinces: site 5 (L5F, signal peptide), site 477 (S477N, RBD), and site 681 (P681H/R, furin cleavage site). Site 681 in particular subtends the S1/S2 boundary that is functionally required for membrane-fusion priming and is the locus of both the P681H mutation diagnostic of Alpha and Omicron and the P681R mutation diagnostic of Delta (Rambaut et al., 2020; O’Toole et al., 2022). Three further sites are recurrent across three provinces (sites 95, 367 and 371), and ten more across two provinces. The full ranked recurrence table is given in Table 2; the spike-positional view is Figure 1a. As Figure 1 makes clear, the recurrent hotspots concentrate in three structurally coherent regions: the N-terminal domain (NTD) “supersite” around positions 18–95, the receptor-binding domain (RBD) saddle (367, 371, 477, 501), and the immediately adjacent S1 C-terminus including D614G and the furin cleavage site.

**Table 2.** Spike codon sites under episodic positive selection in two or more Canadian provinces. EBF range gives the minimum and maximum across all (branch × province) events at that site (capped at 10^12^). Date range is the earliest and latest child-date assigned to any event at that site across provinces; full HPD intervals are provided in Supplementary Table S1.

| Codon | AA change | Domain | Provinces hit | $n$ events | $\log_{10}(\text{EBF})$ range | First event | Last event |
| --- | --- | --- | --- | --- | --- | --- | --- |
| 5 | L5F | Signal peptide | 5 (AB BC ON QC SK) | 70 | 2.0–10.7 | 2020-03-16 | 2021-07-20 |
| 477 | S477N | RBD | 5 (AB BC ON QC SK) | 50 | 2.9–12.0 | 2020-03-28 | 2021-12-22 |
| 681 | P681H/R | Furin cleavage | 5 (AB BC MB ON QC) | 29 | 2.8–9.5 | 2020-04-15 | 2021-08-31 |
| 95 | T95I | NTD | 3 (AB BC QC) | 22 | 2.3–9.7 | 2020-09-01 | 2021-11-29 |
| 367 | V367F | RBD | 3 (AB BC QC) | 9 | 8.9–11.2 | 2020-04-06 | 2021-05-26 |
| 371 | S371F/L | RBD (BA.1) | 3 (AB ON SK) | 4 | 2.9–7.5 | 2021-07-08 | 2022-05-10 |
| 67 | A67V | NTD | 2 (BC QC) | 16 | 7.3–8.9 | 2020-11-07 | 2021-04-27 |
| 18 | L18F | NTD | 2 (BC QC) | 14 | 7.3–9.3 | 2020-08-12 | 2021-04-18 |
| 614 | D614G | S1 C-terminus | 2 (BC QC) | 14 | 4.7–12.0 | 2020-03-13 | 2021-03-06 |
| 1078 | A1078V | HR2 | 2 (BC QC) | 12 | 8.3–$\geq 12$ | 2020-08-11 | 2021-04-10 |
| 75 | D75N | NTD | 2 (BC QC) | 10 | 3.9–8.7 | 2020-03-20 | 2021-05-13 |
| 222 | A222V | NTD | 2 (BC QC) | 10 | $\geq 12$ | 2020-09-18 | 2021-05-18 |
| 501 | N501Y | RBD | 2 (BC QC) | 9 | 4.4–8.1 | 2020-06-07 | 2021-07-24 |
| 20 | T20N | NTD | 2 (AB BC) | 8 | 4.1–8.4 | 2020-12-30 | 2021-05-28 |

**Figure 1.**
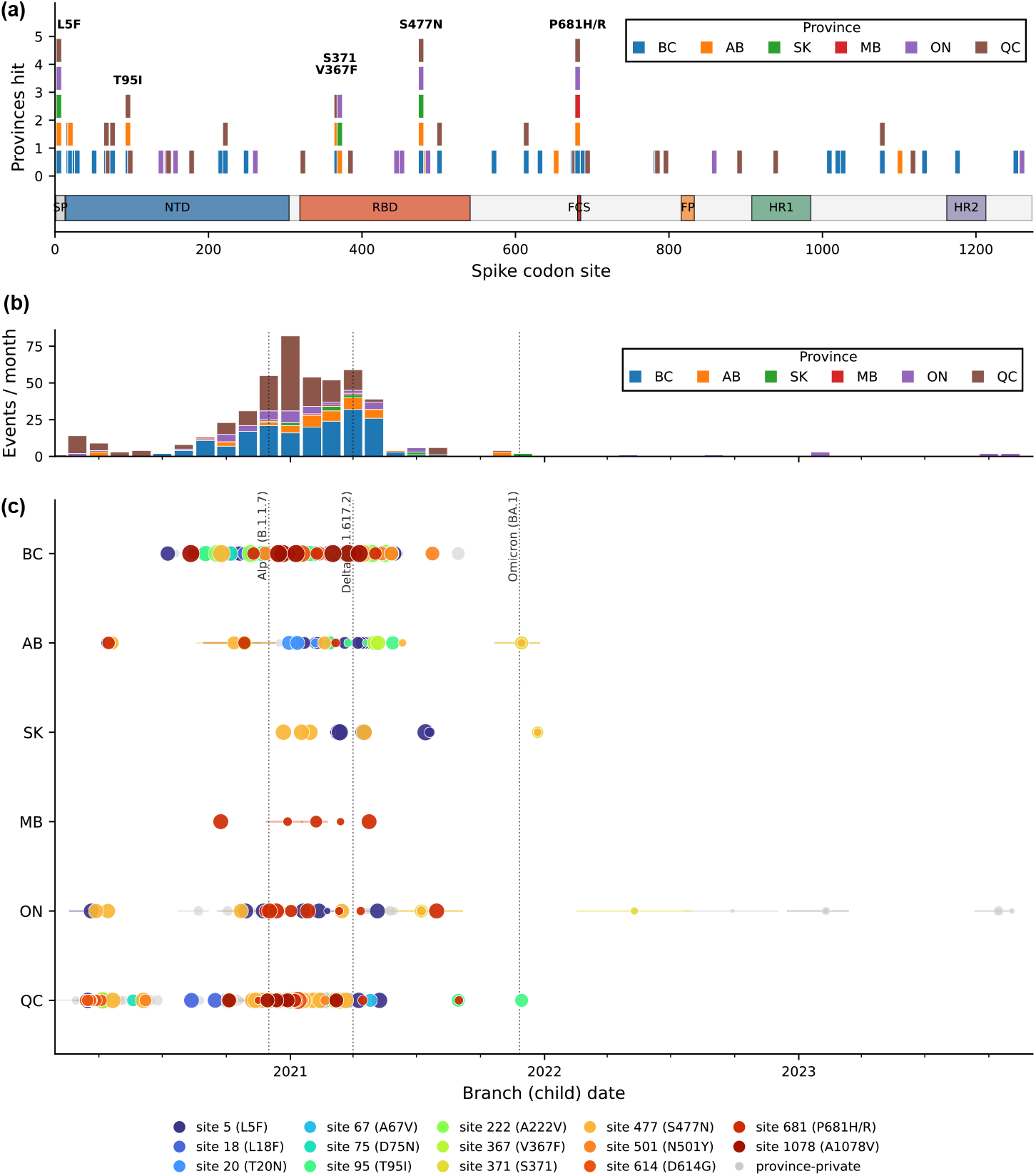
Cross-province recurrence and temporal distribution of episodic positive selection on the SARS-CoV-2 spike protein. **(a)** Codon-site recurrence stack across the entire spike (1,273 codons). Each colored square is a single (province × codon) strict episodic-selection event; columns are stacked by the number of provinces hitting that codon, and tall columns (≥ 3 provinces) carry amino-acid label (see Table 2). The horizontal track underneath shows the domain organization (signal peptide SP, N-terminal domain NTD, receptor-binding domain RBD, furin cleavage site FCS, fusion peptide FP, heptad repeats HR1 and HR2). **(b)** Monthly count of strict events, stacked by province; dotted vertical lines mark the approximate Canadian emergence dates of Alpha (B.1.1.7), Delta (B.1.617.2) and Omicron (BA.1). **(c)** Per-province swimlane: each dot is a single strict episodic-selection event placed at its child-node calendar date on the corresponding BEAST MCC tree; the faint horizontal whisker shows the HPD on the child date. Dots are colored by site for the cross-province (recurrent) hotspots and grey for province-private hits; dot area scales with log_10_(EBF), capped at 10^12^. NB, NS, NL and PE rows are omitted because their BEAST analyses contributed no strict events. See Figure S1 for a detailed view of NTD and RBD/FCS.

### Selection events peak with Alpha and Delta emergence

The temporal distribution of strict events (Figure 1b) is sharply peaked between October 2020 and May 2021, with the four busiest months being January 2021 (82 events), April 2021 (59), December 2020 (55) and February 2021 (54). This window coincides with the Alpha-into-Delta succession in Canada (Figure S2), with each provincial peak displaced by a few weeks consistent with the local epidemic trajectory. The earliest dated event in any province is an event at codon 69 (H69-) in Quebec on 2020-02-28 (HPD 2020-01-28 to 2020-03-17), placing positive-selection signal at the spike before pandemic-era vaccination began anywhere in the world. By contrast, signal at the variant-defining sites 501, 477, 614 and 681 first appears between mid-March and June 2020 in the westernmost provinces; the swimlane in Figure 1c shows that British Columbia’s earliest strict event sits at codon 5 on 2020-07-09. Three Ontario events fall well outside the Alpha-Delta window, in late 2022 and late 2023, all at the BA.1- and Omicron-defining RBD sites 371 and 452; these reflect the longer sampling tail of the Ontario dataset (youngest tip 2023-12-04) and are consistent with continued, low-rate selection on the receptor-binding domain through the Omicron era.

### Vaccine-deployment intensity does not lead a wave of episodic selection

After residualizing both the weekly per-province count of strict selection events and the weekly per-province count of new dose-1 administrations against the national variant-phase covariate, the cross-correlation function (Figure 2) shows no province in which deployment leads a robust burst of selection at a biologically plausible lag (contra the pre-detrended analyses, see SI text). Maximum residualized Pearson correlations fall between *r* = 0.09 and *r* = 0.36, with the lag of maximum correlation distributed across {−4, −2, 0, 0, −4, +13} weeks for {AB, BC, MB, ON, QC, SK}. None of the four provinces with positive maximum-correlation lags falls in the +8 to +24 week window in which a vaccine-driven escape signal would be expected. The 1,000-replicate circular-shift permutation test rejects the null of no association at *α* = 0.05 in only one province (Saskatchewan, perm-*P <* 10^−3^), and there the maximum lag is *negative* (−4 weeks; selection slightly leading vaccine deployment), opposite to the sign expected under an immune-escape hypothesis. British Columbia is borderline (perm-*P* = 0.082) and synchronous (lag 0). The four remaining provinces all yield perm-*P >* 0.17.

**Figure 2.**
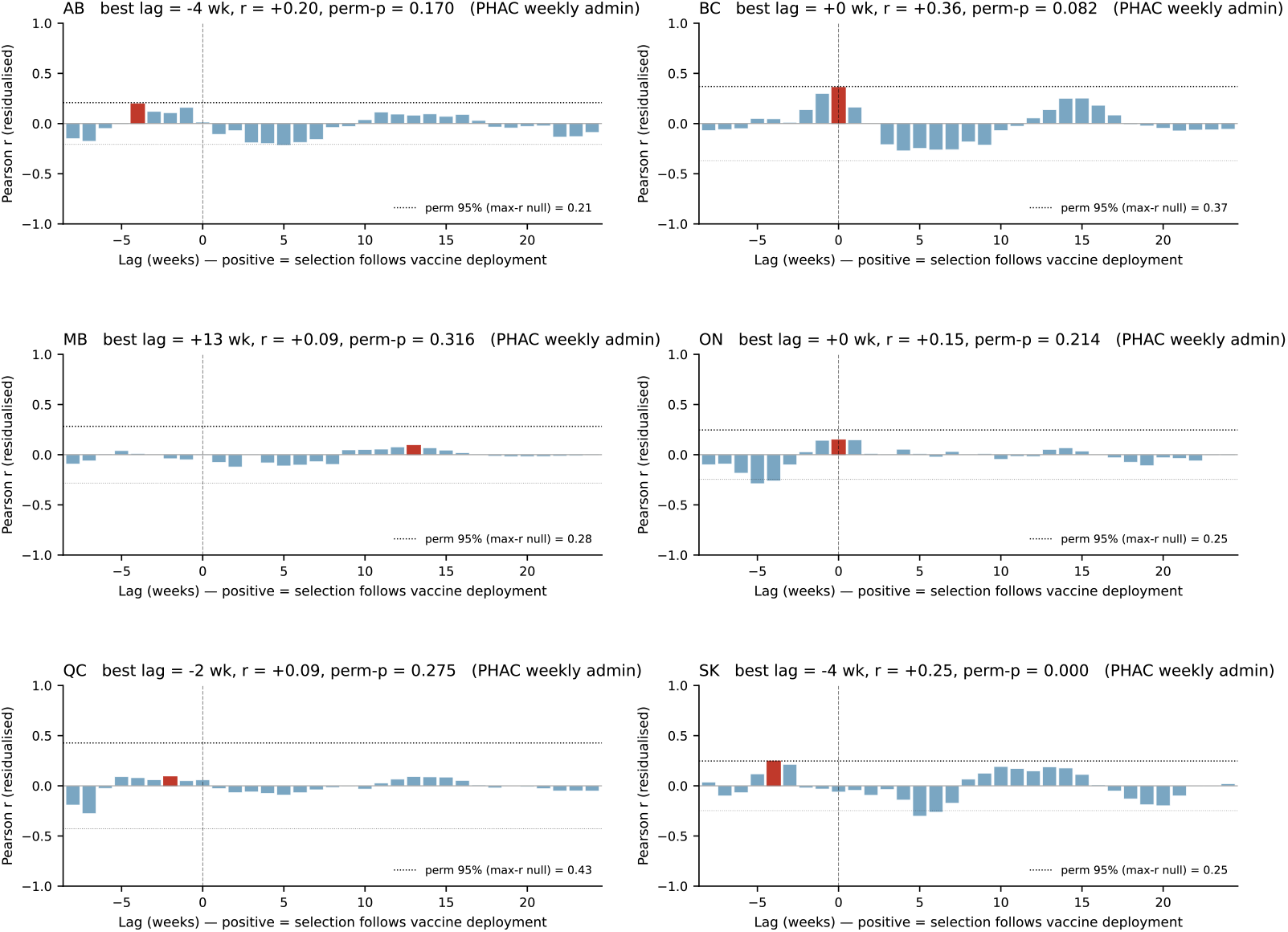
Per-province cross-correlation of the weekly episodic-selection-event count with the weekly count of new dose-1 vaccine administrations, after partialing out the national variant-of-concern phase covariate. For each of the six provinces with ≥5 strict events (AB, BC, MB, ON, QC, SK), the residualized Pearson correlation is plotted at integer weekly lags from −8 to +24 weeks; positive lag means selection follows vaccine deployment. The red bar marks the lag of maximum correlation (reported with its *r* and the empirical *P* -value of a 1,000-replicate circular-shift permutation test on the residualized vaccine series). Dotted lines mark the 95th percentile of the permutation null distribution of the per-replicate maximum *r*. Saskatchewan is the only province whose CCF clears the permutation threshold; its maximum lag is at −4 weeks, opposite the sign expected under a vaccine-driven immune-escape hypothesis.

The CCF figure decisively rules out the impulse-train artifact that afflicted our earlier analyses based on a sparse first-dose-date indicator: with the deployment-intensity series replacing isolated event impulses, the previously-observed cluster of ∼ +3 week peaks across four provinces collapses, and the magnitude of every residualized correlation drops by a factor of approximately 2. We interpret the convergent across-province null finding as evidence that, within the resolution and sample size available, vaccine deployment intensity is not a detectable driver of provincial-scale episodic positive selection on the spike protein. The site-level signal (Figure 1 and Table 2) is most parsimoniously attributed to the rapid rise of variant-defining substitutions during the Alpha-Delta succession, before substantial vaccination-induced immune pressure had accumulated in any province.

## Discussion

### Episodic positive selection on the SARS-CoV-2 spike is dominated by variant-emergence dynamics

Across the eight Canadian provinces with adequate sequence sampling and convergent BEAST inference, episodic positive selection on the spike protein concentrates on a small number of codons that define the biological signature of the major VOC circulating in 2020-2022. The three sites detected as under episodic selection in five of the ten provinces (codons 5, 477 and 681) span the three principal axes of spike adaptation: signal-peptide-proximate N-terminal modulation (L5F), receptor-binding affinity tuning (S477N) and proteolytic priming at the S1/S2 junction (P681H/R) (Wrapp et al., 2020; Walls et al., 2020; Saito et al., 2022; Wrobel et al., 2020). The “three-province” hits at codons 95, 367 and 371 extend the same three-domain pattern: T95I has been a recurrent feature of the NTD antigenic “supersite” (McCallum et al., 2021; McCarthy et al., 2021), V367F lies in the RBD core that contacts ACE2 (Starr et al., 2020), and S371F/L is one of the diagnostic Omicron BA.1 substitutions (Cao et al., 2022; Liu et al., 2022). The “two-province” hits at 18, 20, 67, 75, 222, 501, 614 and 1078 further fill in the same three regions. The functional clustering is consistent with what has emerged from deep mutational scans of the RBD (Greaney et al., 2021; Starr et al., 2020) and from structural analyses of the NTD antigenic supersite (McCallum et al., 2021). It mirrors the consensus that variant-defining substitutions on the spike concentrate at three pressure points: receptor binding, fusion priming, and antibody evasion at the NTD supersite (Carabelli et al., 2023; Markov et al., 2023).

The temporal distribution of these events is concentrated in the late-2020/early-2021 window, peaking January 2021, that corresponds to the Alpha-into-Delta succession in Canada (Berry et al., 2024; Davies et al., 2021; Mlcochova et al., 2021). The earliest dated event in any province (codon 69 in Quebec, 2020-02-28) precedes that wave by nearly a year, suggesting that episodic selection was already operating during the original B.1 era, well before the explicit emergence of named variants. This is fully consistent with the trajectory of D614G itself, whose fitness advantage was established by mid-2020 (Korber et al., 2020; Plante et al., 2020), and which we recover as a two-province hit in British Columbia and Quebec at exactly that date (Table 2, first event at codon 614 on 2020-03-13).

### Spatially replicated emergence is a signature of convergent adaptive evolution

Because each provincial dataset represents a largely independent local epidemic with limited cross-provincial mixing during the relevant sampling windows (Berry et al., 2024), the recurrence of the same codon under episodic selection in five different provinces is most parsimoniously explained by convergent adaptive evolution: the same spike substitutions arose, were selected upward in frequency, and were detected by MEME on independent branches in each provincial tree. This pattern of “spatially replicated parallel evolution” is methodologically the opposite of an introduction-and-spread artifact: if S477N had reached fixation once in eastern Canada and then been introduced into successive provinces, we would expect a single selection event near the root of the introduction subtree in each province, not the dozens of branch-level hits we recover at codon 477 in five provinces. The within-province distribution of dates at these recurrent sites (Table 2 and Figure 1c) further supports parallel selection: site 477 events span 2020-03 to 2021-12 across five provinces, and site 681 events span 2020-04 to 2021-08 across five provinces, far broader temporal windows than would result from a single origin.

We note that the whole-gene BUSTED test was non-significant in every province (Ontario at *P*_*BUSTED*_ = 0.051 being the only borderline case). This combination (gene-wide null, narrow site-level signal) is exactly the pattern expected when selection is intense but restricted to a small fraction of codons, and it is consistent with the general observation that the *dN/dS* regime of SARS-CoV-2 across the entire spike is close to neutrality, with the adaptive signal concentrated on a handful of functionally critical residues (Murrell et al., 2015; Carabelli et al., 2023; Markov et al., 2023).

### Vaccine-deployment intensity is not a detectable driver of provincial selection

After partialling out the national variant-of-concern phase covariate, weekly per-province dose-1 administration intensity does not systematically lead a wave of episodic selection at any biologically plausible lag (Figure 2). The maximum residualized correlations are modest (*r* = 0.09-0.36), the maximum-correlation lag is distributed across {−4, −2, 0, 0, −4, +13} weeks, and only one province (Saskatchewan) clears a circular-shift permutation test at *α* = 0.05. All this while there is a negative lag, with selection slightly preceding vaccine deployment, opposite to the sign expected under a vaccine-driven escape hypothesis. This suggests that, at provincial scale, with weekly resolution, after detrending the variant-emergence cycle, vaccine deployment intensity is not a detectable driver of episodic positive selection at the SARS-CoV-2 spike in this dataset.

This negative finding is consistent with the broader literature. The major immune-escape sweeps detected on the spike during the period covered by our analysis (D614G, the Alpha RBD/NTD/FCS mosaic, and the Delta L452R/T478K/P681R mosaic) arose either before the start of vaccination or well before population-level vaccine-induced immunity could plausibly have generated provincial selection pressure (Korber et al., 2020; Davies et al., 2021; Mlcochova et al., 2021; Carabelli et al., 2023). Convergent neutralizing-antibody escape under vaccine-induced immunity has been demonstrated experimentally (Greaney et al., 2021; Liu et al., 2022; Cao et al., 2022), but the magnitude of that selection on circulating populations during early- and mid-2021 in Canada was likely small relative to the much stronger fitness gains delivered by Alpha-then-Delta variant emergence (Markov et al., 2023; Carabelli et al., 2023). The picture would likely be different if the analysis had been extended into the BA.5 and XBB era, when waning vaccine-induced immunity and recurrent boosting created a more sustained antibody-pressure landscape, but our provincial sample sizes in that period (especially outside Ontario) are too thin to support the same analysis.

### Methodological considerations

Three methodological choices in this work deserve explicit comment. First, we date selection events by mapping each MEME-significant (branch × site) onto a BEAST-time-calibrated phylogeny via descendant tip-set matching, rather than by re-running MEME with a fixed BEAST topology. The latter would be more elegant but is computationally infeasible at the per-province sample sizes considered here, and the tip-set approach achieved 94–97% branch matching across the converged provinces, which is a reasonable trade-off.

Second, we used a 1,000-replicate circular-shift permutation null to assess significance of the per-province cross-correlation peaks. This is a deliberately conservative choice as circular shifts preserve the marginal distribution and autocorrelation of the vaccine series, while randomizing its phase relative to the selection series, so the null is internally consistent with the smoothness of the data . However, it is also somewhat under-powered when the test statistic (maximum *r* across lags) has a tight null distribution because of limited time-series length. The corollary is that our perm-*P* values, especially the borderline-significant British Columbia result (perm-*P* = 0.082), should be treated as evidence rather than proof.

Third, we constructed the variant-phase covariate at the national level rather than the provincial level. The Canadian VOC succession was sufficiently synchronous across provinces that a national indicator captures most of the variance, but Western and Atlantic provinces did differ by ∼ 2 weeks in the timing of the Alpha-Delta transition (Berry et al., 2024). A provincial phase covariate (once sub-provincial CoVariants-Canada or wastewater data are reachable in analysis-ready form) would tighten the residualization and could sharpen the cross-correlation results, particularly at the boundary between phases.

Three limitations of the current analysis are worth noting. The first is the sampling-effort confound. Episodic-selection events are detected on branches of trees that are themselves built from a sample of genomes; if genome sequencing intensity is correlated with both case-wave intensity and vaccine-campaign intensity (which was the case in Canada through 2020–2022), then the residualized cross-correlation analysis cannot fully distinguish “vaccine deployment selects for spike substitutions” from “vaccine deployment correlates with the surveillance-attention regime that generates the data we analyze.” Including a per-province sequencing denominator (genomes sequenced per week) as an additional covariate in the partialling step would be a straightforward extension, once sub-provincial sequencing-effort time series are released.

The second is the strictness of our MEME thresholds. We required *P*_*MEME*_ ≤ 0.10 *and* EBF ≥ 100 for an event to be called strict. These thresholds are conservative and select for high-confidence, branch-level episodic events, but they will miss the kind of broadly-distributed, low-amplitude selection signal that a generalized vaccine-induced antibody pressure might impose. Repeating the analysis at the suggestive threshold (*P*_*MEME*_ ≤ 0.20) recovers 812 events with the same qualitative pattern and the same null cross-correlation result, suggesting that the conclusions are robust to threshold choice within the regime explored, but a complementary analysis based on FUBAR or aBSREL site-level posteriors would be a useful sanity check (Kosakovsky Pond et al., 2020).

The third is the small-sample regime in four provinces (New Brunswick, Newfound-land & Labrador, Nova Scotia, and Prince Edward Island). Our pipeline correctly flags Prince Edward Island as non-converged and excludes it from interpretation; New Brunswick, Newfoundland & Labrador and Nova Scotia all mix acceptably but their wide root-height HPDs (Table 1 of Results) reflect their small sample sizes (∼29, ∼21 and ∼67 tips, respectively) and they contribute no strict episodic-selection events. A future extension that reanalyses these alignments under a more informative prior on the clock rate (a Brownian-bridge prior anchored on the global SARS-CoV-2 substitution-rate posterior) might allow rate inference to piggy-back on the global signal where the provincial signal is weak.

Two natural extensions of the present work are worth highlighting. First, integrating the per-event EBFs and dates into a phylodynamic “selection landscape” (a per-codon, per-week selection coefficient inferred jointly from the time-calibrated tree and the MEME posteriors) would replace the current event-counting analysis with a continuous intensity field, increasing statistical power and removing the threshold dependence. Second, layering on natural-immunity exposure (seroprevalence) in addition to vaccine deployment would allow the relative contributions of infection-induced and vaccine-induced selective pressure to be separated, which is the question the present analysis really wants to address.

### Conclusions

At the resolution of provincial Canadian SARS-CoV-2 datasets, episodic positive selection on the spike protein is all about variant emergence, not about vaccine deployment. The same handful of functionally critical codons (the NTD antigenic supersite, the RBD/ACE2 contact saddle and the S1/S2 furin-cleavage region) is struck by independent positive-selection events in province after province, on a calendar window that aligns with Alpha-then-Delta emergence rather than with the rollout of any specific vaccine product. After detrending against a national variant-phase covariate and using actual weekly per-province dose-1 administration counts, we find no evidence in our dataset that vaccine-deployment intensity drives a lagged wave of episodic spike selection at provincial scale. The broader question of whether population-scale vaccine-induced antibody pressure measurably shapes the evolution of circulating respiratory viruses at the level of detectable site-by-branch selection events, remains worth pursuing with finer-grained sampling, exposure data and selection-landscape methods, of which this work is a starting framework.

## Acknowledgements

We gratefully acknowledge all data contributors, i.e., the Authors and their Originating laboratories responsible for obtaining the specimens, and their Submitting laboratories for generating the genetic sequence and metadata and sharing via the GISAID Initiative, on which this research is based. We would also like to thank the Digital Research Alliance of Canada (DRAC), formerly known as Compute Canada, for compute time.

## Data availability

The code developed for this work is available from https://github.com/sarisbro/data.

