## Supplementary material for "Variant emergence, not vaccine deployment, drives episodic positive selection on the SARS-CoV-2 spike at provincial scale in Canada": SI

This document contains the supplementary tables, figures, and reproducibility appendix that accompany the manuscript on episodic positive selection on the SARS-CoV-2 spike protein across Canadian provinces. Section 1 reports per-province dataset sizes and the rate at which HyPhy branches were matched to homologous BEAST branches by descendant tip-set. Section 2 reports per-parameter convergence diagnostics for the two-chain BEAST runs. Section 3 defines the national variant-phase covariate used to detrend the cross-correlation analysis. Section 4 explains the importance of detrending the data. Section 5 gives the derivation of the empirical Bayes factor used to filter (branch $\times$ site) selection events. Section 6 contains three supplementary figures: an alternative double-column rendering of the main events figure that zooms into the NTD and RBD/FCS; the pre-detrending monthly cross-correlation analysis that motivated the move to weekly resolution and variant-phase residualization; and the provincial vaccine-rollout time series that underlies the deployment side of the detrended cross-correlation analysis. Section 7 lists the analysis scripts.

#### 1 Provincial dataset sizes and tree-matching

Per-province sample sizes after redundancy collapse, and the rate at which HyPhy branches matched a homologous BEAST branch via descendant tip-set (Materials and Methods, *Dating selection events*), are reported in Table S1. Match rates exceed 93% in every province except Nova Scotia, where small numbers of short internal branches near the basal cluster contributed disproportionate topological discordance.

**Table S1. Per-province dataset sizes and HyPhy/BEAST branch-matching rates.** The cleaned codon alignment for each province was reduced to non-redundant sequences at 100% nucleotide identity (CD-HIT-EST; Fu et al., 2012); $n$ is the number of tip sequences retained. The HyPhy maximum-likelihood phylogeny and the BEAST MAP tree were built independently from the same alignment; *branches matched* reports the number of HyPhy branches whose descendant tip-set is identical to that of a unique BEAST branch.

| Province | $n$ tips | HyPhy branches | Branches matched | Match rate (%) |
|---|---|---|---|---|
| Alberta (AB) | 576 | 1,151 | 1,087 | 94.4 |
| British Columbia (BC) | 1330 | 2,658 | 2,510 | 94.4 |
| Manitoba (MB) | 152 | 303 | 286 | 94.4 |
| New Brunswick (NB) | 29 | 57 | 56 | 98.2 |
| Newfoundland & Labrador (NL) | 21 | 40 | 37 | 92.5 |
| Nova Scotia (NS) | 67 | 133 | 119 | 89.5 |
| Ontario (ON) | 588 | 1,175 | 1,105 | 94.0 |
| Prince Edward Island (PE) † | 10 | 19 | 18 | 94.7 |
| Quebec (QC) | 1121 | 2,240 | 2,101 | 93.8 |
| Saskatchewan (SK) | 168 | 335 | 313 | 93.4 |

† Prince Edward Island is shown for completeness but its BEAST analysis did not converge (see Section 2); event dates derived from this run are not interpretable.

#### 2 Per-province MCMC convergence diagnostics

Effective sample size (ESS, computed via Geyer's initial monotone sequence estimator on the combined post-burn-in chain; Geyer, 1992) and the Gelman-Rubin potential scale-reduction factor ($\hat{R}$, computed between the two independent chains; Gelman and Rubin, 1992) are reported for the six continuous parameters that drive the molecular-clock fit (Table S2). The acceptance criteria stated in the main Methods (ESS $\geq$ 200 and $\hat{R} \leq 1.05$) are met for almost all parameter-province pairs. The conspicuous exceptions are: (i) Quebec's tree-length and constant-population-size parameters, both of which mix poorly because of the extreme node-density of the QC posterior MAP tree; (ii) British Columbia's clock-rate and constant-popSize parameters, which have $\hat{R} > 1.05$ but ESS far above the threshold and pose only minor concern; (iii) Prince Edward Island, whose root-height and total-tree-length posteriors are biologically meaningless (means of $\sim 10^6$ years), the chain technically samples the prior because the tip number is too small to constrain the coalescent; PE results are not interpreted in the main text.

**Table S2. Per-province MCMC convergence diagnostics for the six continuous BEAST parameters.** Each cell reports the combined-chain ESS over the Gelman-Rubin $\hat{R}_{\text{run1 vs run2}}$ statistic. Bold font marks parameter values that fail at least one of the two acceptance criteria (ESS $\geq$ 200, $\hat{R} \leq 1.05$). PE failed on multiple parameters and on the joint likelihood; values not shown.

| Province | clock.rate | rootHeight | kappa | alpha | popSize | treeLength |
|---|---|---|---|---|---|---|
| AB | **60 / 1.010** | **19 / 1.067** | 15,667 / 1.000 | 18,807 / 1.000 | **47 / 1.012** | **32 / 1.020** |
| BC | 35 / **1.141** | 3,469 / 1.000 | 2,088 / 1.008 | 6,756 / 1.008 | 48 / **1.116** | **39 / 1.144** |
| MB | **66 / 1.004** | 716 / 1.000 | 7,491 / 1.000 | 18,049 / 1.000 | **57 / 1.004** | **51 / 1.004** |
| NB | 2,698 / 1.000 | 2,892 / 1.000 | 19,686 / 1.000 | 18,651 / 1.000 | 2,525 / 1.000 | 2,317 / 1.000 |
| NL | 2,778 / 1.000 | 4,080 / 1.000 | 18,317 / 1.000 | 16,094 / 1.000 | 4,389 / 1.000 | 3,762 / 1.000 |
| NS | 1,074 / 1.000 | 1,875 / 1.000 | 5,317 / 1.000 | 18,068 / 1.000 | 1,315 / 1.000 | 1,098 / 1.000 |
| ON | **142 / 1.000** | **117 / 1.010** | 383 / 1.001 | 10,643 / 1.000 | **85 / 1.000** | **74 / 1.001** |
| PE | — | — | 19,313 / 1.000 | 17,498 / 1.000 | — | — |
| QC | 23 / **1.252** | 309 / 1.034 | 1,545 / 1.005 | 18,682 / 1.003 | **16 / 1.353** | **14 / 1.419** |
| SK | **176 / 1.004** | 1,010 / 1.003 | 15,967 / 1.000 | 19,325 / 1.000 | **169 / 1.005** | **142 / 1.007** |

#### 3 National variant-of-concern phase definition

The variant-phase covariate used to partial out variant-emergence dynamics before the cross-correlation analysis (Materials and Methods, *Variant-phase covariate*) is defined in Table S3. Phase boundaries are the calendar week in which the focal lineage first exceeded 50% of sequenced isolates in the integrated PHAC clinical and wastewater surveillance system (Berry et al., 2024), with cross-checks against the National Collaborating Centre for Infectious Diseases variant updates (National Collaborating Centre for Infectious Diseases, 2024). Each weekly time-step of the analysis was assigned exactly one of the nine phases.

#### 4 On the importance of detrending

The first version of our cross-correlation analysis represented the provincial vaccine signal as a Gaussian-smoothed monthly impulse train of the curated first-dose dates (Section 3; Table S3), and compared it against the monthly per-province count of strict episodic-selection events. The resulting per-province cross-correlation functions (Figure S3) suggest that each of the six provinces shows a single broad peak with maximum correlation Pearson $r \in [0.60, 0.77]$ and best lag clustered at $\{-1, 0, 0, +1, +1, +1\}$ months, and four of six panels rise above the 95% circular-shift permuta-

**Table S3. National variant-of-concern phase boundaries used for residualization.** Phase *B.1 wildtype* acts as the reference category; *post-BA.5* groups the BQ.1 and related Omicron sublineages that briefly co-circulated before XBB became dominant.

| Phase label | First dominant date | Last dominant date | Weeks in window |
|---|---|---|---|
| `B1_wildtype` | 2020-01 (start) | 2021-02-28 | 64 |
| `alpha` (B.1.1.7) | 2021-03-01 | 2021-06-30 | 18 |
| `delta` (B.1.617.2) | 2021-07-01 | 2021-12-14 | 24 |
| `omicron_BA1` | 2021-12-15 | 2022-02-28 | 11 |
| `omicron_BA2` | 2022-03-01 | 2022-06-30 | 17 |
| `omicron_BA5` | 2022-07-01 | 2022-10-31 | 18 |
| `omicron_postBA5` | 2022-11-01 | 2023-03-31 | 21 |
| `XBB` | 2023-04-01 | 2023-10-31 | 31 |
| `JN1` | 2023-11-01 | 2024-01-22 (end) | 12 |

tion threshold. Read at face value, this would suggest strong evidence that vaccine deployment is synchronously coupled to provincial spike selection.

That reading is, however, almost entirely an artefact of the analytical setup. Both time series share the same wave-like temporal envelope (variant-of-concern emergence and the surveillance intensity that accompanies it), and any two co-trending series will produce large Pearson correlations under classical spurious-regression conditions (Yule, 1926; Granger and Newbold, 1974). The high $r$ values in Figure S3 therefore confirm only that the mass-vaccination campaign occupied the same calendar window as the Alpha-into-Delta selection peak (Figure 1b of the main text), not that the two are causally coupled. Moving the analysis to weekly resolution and partialling out the nine-phase national variant-of-concern covariate (Section 3; Materials and Methods, *Detrended cross-correlation analysis*) before computing the cross-correlation removes this shared-trend variance from both series and re-asks the question on the within-variant-phase residuals, the quantity that maps onto an immune-pressure mechanism rather than onto temporal coincidence. The result of the corrected analysis (Figure 2 of the main text) is markedly different: maximum residualized correlations fall to $r \in [0.09, 0.36]$, the lag of maximum correlation scatters across $\{-4, 0, +13, 0, -2, -4\}$ weeks instead of clustering at zero, and only Saskatchewan clears the permutation threshold, but at $-4$ weeks, opposite the sign expected for a vaccine-driven immune-escape pathway.

### 5 Empirical Bayes factor for branch-level events

For a given site $i$, MEME fits a two-class $\omega$ mixture and returns the per-branch posterior probability that branch $j$ belongs to the positive-selection class, $\Pr(\omega_+|j, i)$, and the estimated mixing weight of that class at the site, $p_i^{\text{pos}}$. The latter is interpretable as the prior probability that any given branch belongs to the positive-selection class *before observing the branch-level data.* The branch-level empirical Bayes factor is the ratio of posterior odds to prior odds:

$$\text{EBF}_{ij} \;=\; \frac{\Pr(\omega_+ \mid j, i)}{1 - \Pr(\omega_+ \mid j, i)} \Bigg/ \frac{p_i^{\text{pos}}}{1 - p_i^{\text{pos}}}. \tag{S1}$$

EBF is exactly 1 when the branch's posterior matches the site's prior (no branch-specific evidence) and rises rapidly when the branch-level data make the positive-selection class more likely than average for that site. Equation (S1) is the same statistic proposed by Murrell et al. (2012) as part of the original MEME framework. Throughout this work we filter (site × branch) events on the joint criterion $P_{MEME} \leq 0.10$ and EBF $\geq 100$ (strict), or $P_{MEME} \leq 0.20$ and EBF $\geq 100$ (suggestive); the EBF threshold of 100 corresponds to “decisive” evidence on the Jeffreys-Kass-Raftery scale.

### 6 Supplementary figures

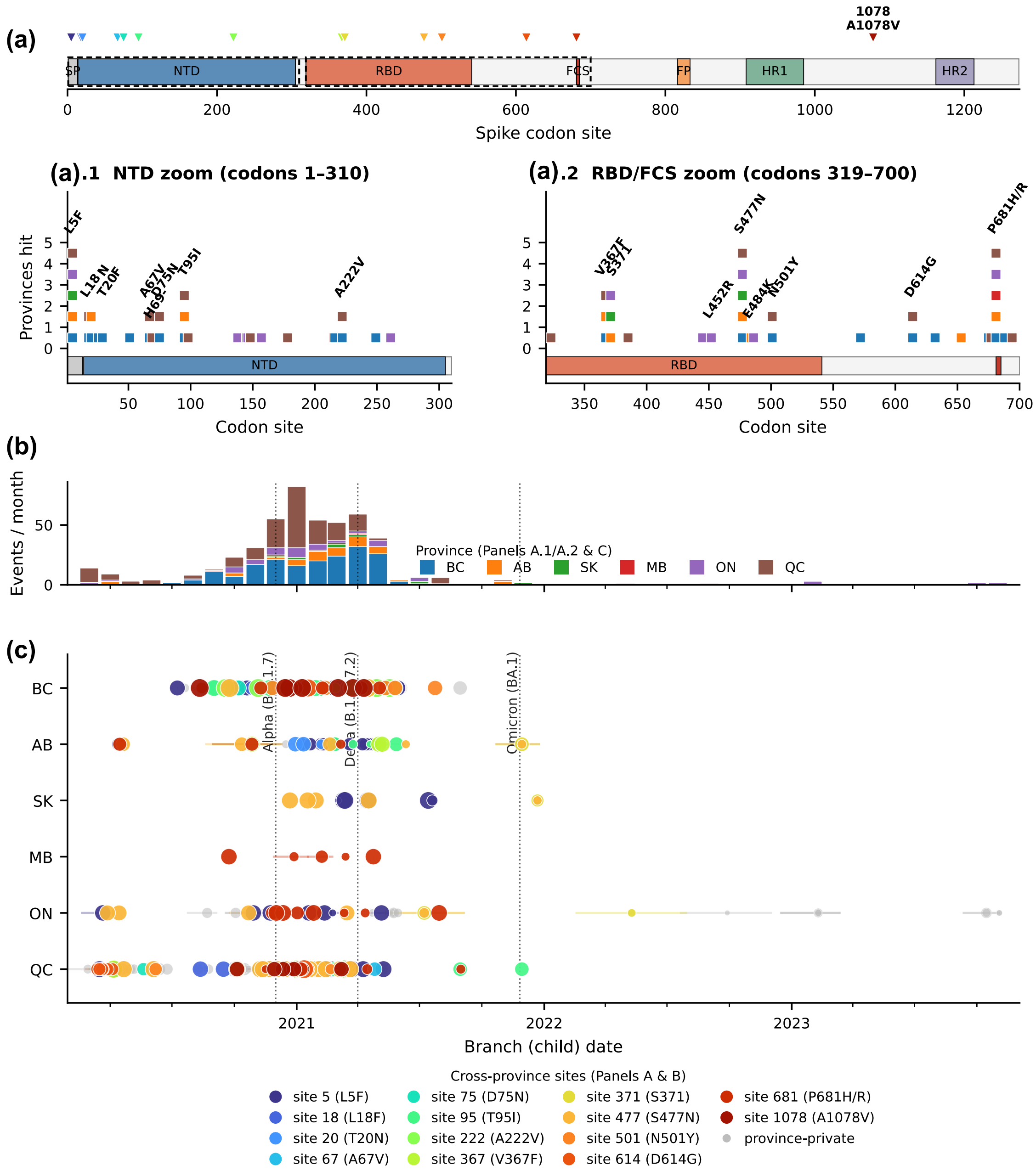


**Figure S1. Alternative rendering of the main events figure with explicit NTD and RBD/FCS zoom panels.** **(a)** (top strip) shows the full-spike domain track with each cross-province ($\geq$ 2-province) site marked as a colored triangle; the dashed brackets indicate the two zoom regions detailed below. **(a.1)** (NTD zoom, codons 1-310) and **(a.2)** (RBD/FCS zoom, codons 319-700) show the recurrence stack for sites in those domains, with diagonally-rotated amino-acid labels above each column. **(b)** and **(c)** are identical to those of Figure 1 in the main text. **(a)** overview labels site 1078 (A1078V), the only recurrent site falling outside the two zoom windows.

The provincial vaccine rollout that supplies the deployment side of the cross-correlation analysis (Materials and Methods, *Detrended cross-correlation analysis*) is shown in Figure S2. Daily

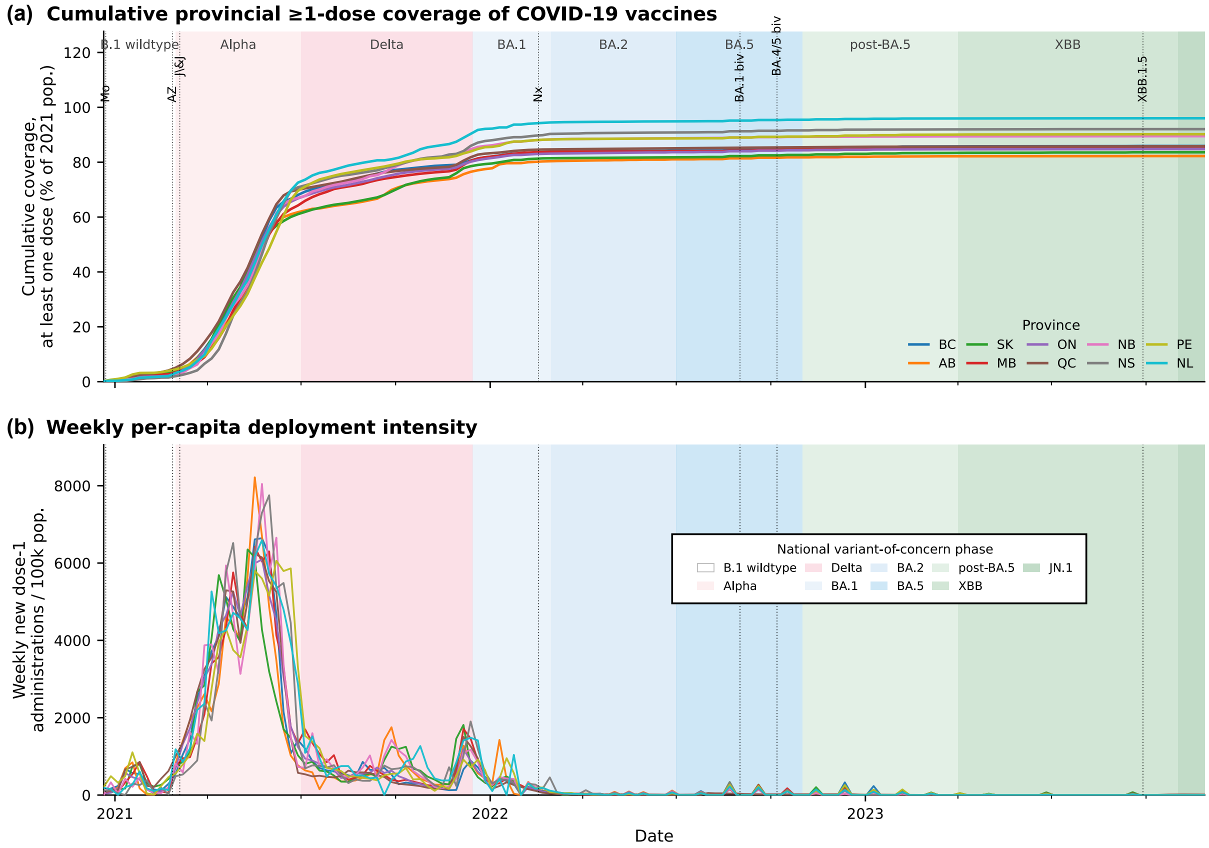


**Figure S2. Provincial COVID-19 vaccine rollout in Canada, 2020-2023. (a)** Cumulative one-dose coverage per province, expressed as a percentage of the 2021 mid-year provincial population estimate (Statistics Canada, 2021). Each line is one province; province colors match the palette used elsewhere in the manuscript. Thin vertical dotted lines mark the Health Canada authorization dates of the principal vaccine products, with abbreviated labels staggered to avoid horizontal overlap: *Pf* = Pfizer-BioNTech BNT162b2, *Mo* = Moderna mRNA-1273, *AZ* = AstraZeneca/Covishield ChAdOx1-S, *J&J* = Janssen Ad26.COV2.S, *Nx* = Novavax NVX-CoV2373, *BA.1 biv* = Moderna mRNA-1273.214 (original + BA.1) bivalent booster, *BA.4/5 biv* = Pfizer and Moderna BA.4/5-adapted bivalent boosters, *XBB.1.5* = monovalent XBB.1.5-adapted boosters. **(b)** Weekly new dose-1 administrations per 100,000 population per province, on the same calendar axis as in **(a)**. The mass-vaccination campaign of mid-2021 dominates the figure; the subsequent small peaks correspond to the BA.1 bivalent (autumn 2022), BA.4/5 bivalent (autumn 2022) and XBB.1.5 monovalent (autumn 2023) booster campaigns. **Background shading** marks the nine national variant-of-concern phases used to residualize both the selection and deployment series before the cross-correlation analysis of Figure 2 (Section 3; Table S3).

per-province dose-1 administration counts from the COVID-19 Canada Open Data Working Group release (Berry et al., 2023) were clipped at zero, aggregated to Monday-anchored weekly sums, and normalized to the 2021 mid-year provincial population estimates of Statistics Canada (Statistics Canada, 2021). Reading the two panels horizontally makes clear that the bulk of the mass-vaccination campaign was concentrated in the Alpha-into-Delta window of 2021-Q2/Q3, the same calendar window in which the strict episodic-selection events of Figure 1 are most numerous. This temporal coincidence is the source of the confounding that the variant-phase residualization step (Section 3) is designed to address.

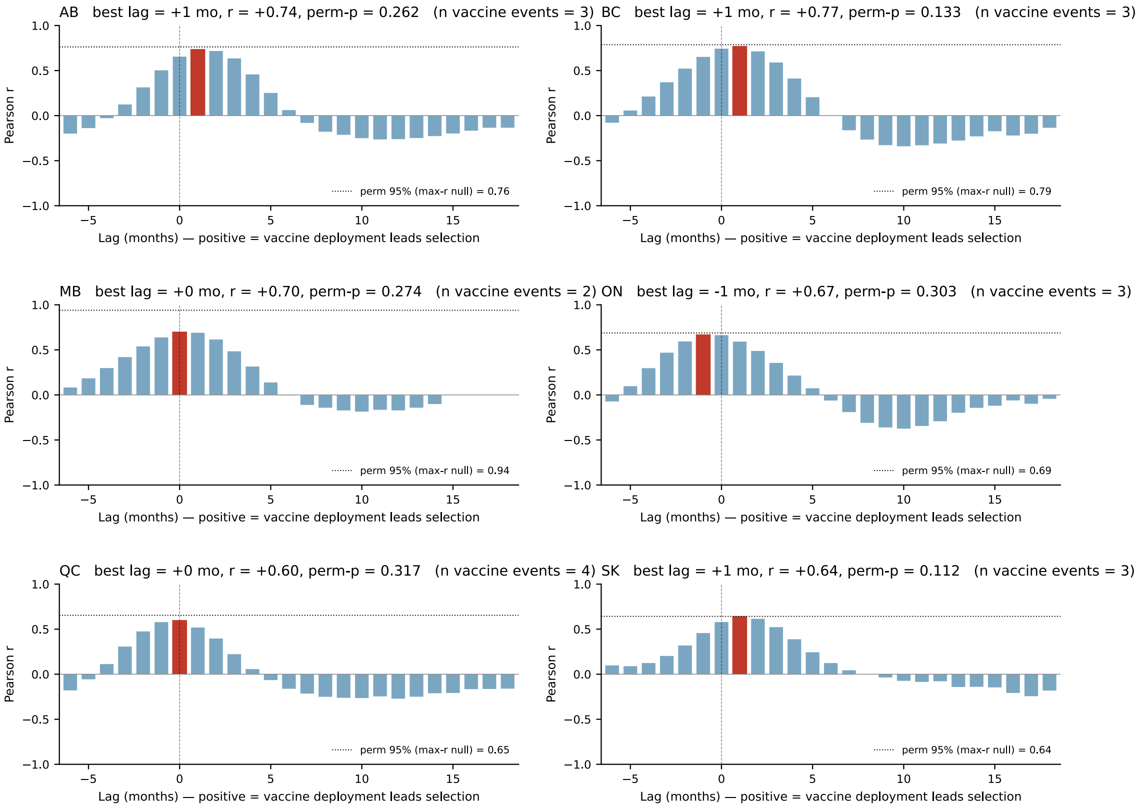


**Figure S3. Pre-detrending monthly cross-correlation analysis (six provinces).** Per-province Pearson correlation between the monthly count of strict episodic-selection events and a Gaussian-smoothed monthly impulse train of provincial first-dose dates (Materials and Methods, *Detrended cross-correlation analysis*), at lags from $-6$ to $+18$ months (positive lag = vaccine deployment leads selection). Red bar marks the lag of maximum correlation. Dotted line is the 95th percentile of the per-replicate maximum-$r$ distribution from a 1,000-replicate circular-shift permutation null. The pattern of synchronous-or-near peaks across all six provinces (best lag $\in \{-1, 0, 0, +1, +1, +1\}$ months, $r \in [0.60, 0.77]$) was the motivation for moving to weekly resolution and partialling out the national variant-phase covariate (Figure 2 of the main text); after detrending, the per-province peaks scatter and the apparent across-province coherence is lost.

### 7 Software and data availability

All analyses are reproducible from the scripts archived in the repository at `https://github.com/sarisbro/data`:

- `0.clean_alignment.py`: forked implementation of trimAl.
- `1.split_by_province.R`: split the alignment of the S gene by province.
- `2.generate_hyphy_scripts.R`: generate the HyPhy scripts for the MEME and BUSTED analyses.
- `3.distribute.hyphy.sh` and `4.run_hyphy.sh`: dispatch the HyPhy runs (local and SLURM versions).
- `5.check_convergence.py`: ESS / $\hat{R}$ diagnostics producing `convergence_summary.tsv` and `convergence_flags.tsv`.

- `6.compare_provinces.py`: cross-provincial posterior extraction producing the `compare_provinces.xlsx` workbook.
- `7.run_combine_annotate.sh`: per-province BEAST chain combination and MAP tree summarization, with parallel dispatch via `wait -n`.
- `8.date_selection_events.py`: branch-matching by tip-set normalization, EBF computation, and assignment of parent/child calendar dates and HPD intervals to each (site × branch) event. Produces `selection_events.tsv`, `selection_events_suggestive.tsv`, and `selection_events_summary.x`
- `9.clean_codon_alignment.py`: column-consensus imputation of IUPAC ambiguities and stop-codon replacement, with provenance log.
- `10.plot_selection_events.py` and `11.plot_selection_events_fullwidth.py`: code for the two publication-format renderings of the main events figure.
- `12.build_vaccine_timeline.py`: curated vaccine-timeline CSV builder.
- `13.correlate_selection_vaccines.py` (monthly) and `14.correlate_selection_vaccines_weekly.py` (weekly, phase-detrended). The weekly script auto-detects the `vaccine_administration_dose_1_pt.csv` drop-in from the COVID-19 Canada Open Data Working Group (Berry et al., 2023) and falls back to the curated first-dose impulse train when the drop-inis not present.
